# Proteogenomic analysis of aneuploidy reveals divergent types of gene expression regulation across cellular pathways

**DOI:** 10.1101/2021.12.07.471176

**Authors:** Pan Cheng, Xin Zhao, Lizabeth Katsnelson, Raquel Moya, Jasmine Shwetar, David Fenyö, Teresa Davoli

## Abstract

How cells control gene expression is a fundamental question. The relative contribution of protein-level and transcript-level regulation to this process remains unclear. Here we perform a proteogenomic analysis of tumors and untransformed cells containing somatic copy number alterations (SCNAs). By revealing how cells regulate transcript and protein abundances of SCNA-containing genes, we provide insights into the rules of gene regulation. While gene compensation mainly occurs at the protein level across tumor types, genes gained or lost show surprisingly low protein compensation in lung and high RNA compensation in colon cancer. Protein complex genes have a strong protein-level regulation while non-complex genes have a strong transcript-level regulation. Exceptions are plasma membrane protein complexes showing a very low protein-level regulation. Strikingly, we find a strong negative association between the degree of transcript-level and protein-level regulation across genes and pathways. Moreover, genes participating in the same pathway show similar degree of transcript- and protein-level regulation. Pathways including translation, splicing and mitochondrial function show a stronger protein-level regulation while cell adhesion and migration pathways show a stronger transcript-level regulation. These results suggest that the evolution of gene regulation is shaped by functional constraints and that many cellular pathways tend to evolve a predominant mechanism of gene regulation, possibly due to energetic constraints.

**Highlights:**

- Proteogenomic analyses of cancer SCNAs reveal tissue specificity in gene compensation.
- Genes gained or lost show surprisingly low protein compensation in lung cancer and unexpected RNA compensation in colon cancer.
- We use DNA-RNA and RNA-protein correlations to infer the degree of RNA-level and protein-level regulation.
- Protein complex genes and non-complex genes show high protein-level and RNA-level regulation, respectively.
- Plasma membrane complexes are an exception showing more RNA-level than protein-level regulation than other complex genes.
- Genes participating in the same pathway show similar degree of RNA-level and protein-level regulation.
- There is a strong negative relationship between the RNA- and protein-level regulation among pathways, suggesting that they are regulated either at the protein or at the RNA level.
- Genes involved in RNA processing and protein synthesis are upregulated in highly aneuploid tumors, especially at the protein level.

## Introduction

The expression level of each gene depends on the regulation of its transcript abundance (RNA-level regulation) and of its protein abundance (protein-level regulation) through synthesis, processing and degradation of its transcript and protein, respectively. RNA-level and protein-level regulation is tightly controlled not only to adapt to changes in environmental conditions, but also as a mechanism to optimize energy consumption (Franks et al., 2017; Wagner, 2005). Certain genes are thought to have a predominant mechanism of regulation, either at the RNA level or at the protein level. For example, *HTERT*, encoding human telomerase, has a strong RNA-level regulation through transcriptional and splicing control (Cong et al., 2002; Lazzerini-Denchi & Sfeir, 2016). In contrast, *GCN4* (and its homolog *ATF4*) has a strong protein-level regulation through increased translation under ER (endoplasmic reticulum) stress (Holcik & Sonenberg, 2005). In addition, cell cycle genes such as *CDT1* and *CDC25A* are strongly regulated at the protein level through protein degradation (Emanuele et al., 2011). The relative contribution of RNA-level and protein-level regulation to control the expression level of human genes is currently incompletely understood.

To assess gene regulation, several studies have investigated the RNA and protein half-lives or the association between RNA and protein abundance across human genes in cells or tissues (Gygi et al., 1999; Marguerat et al., 2012; Mathieson et al., 2018; McShane et al., 2016; Schwanhäusser et al., 2011). Another way to investigate gene regulation is to measure how RNA and protein abundances change upon alterations in DNA copy number of a given gene or SCNAs (somatic copy number alterations; amplifications and deletions) that naturally occur in human cancers or that are experimentally engineered in cells. Previous proteomics analyses of aneuploid yeast and human cells have shown that, while most genes exhibit high correlation between DNA copy number and RNA abundance, a significant fraction of genes (20-30%) do not show protein abundance changes that are proportional to the DNA or RNA changes (Gonçalves et al., 2017; Jovanovic et al., 2015; McShane et al., 2016; Stingele et al., 2012; Torres et al., 2007). In other words, the protein abundance is attenuated with respect to what it is expected based on its DNA change (gene compensation). In particular, genes whose products are participating in protein complexes (protein complex genes) show a stronger compensation than genes that are not part of complexes (non-complex genes). Importantly, a recent study showed that the genes showing compensation at the protein level in aneuploid cells have also a high degree of protein-level regulation in normal diploid cells (McShane et al., 2016). In other words, this study found that protein compensation in aneuploid cells is associated with protein regulation (degradation patterns) in normal cells. This supports the fact that studying how protein levels are affected by SCNAs in (aneuploid) cancer cells can inform us on how genes are regulated in normal (non-aneuploid) cells (Taggart et al., 2020).

Although such studies have advanced our understanding of how cells regulate RNA and protein abundances of genes that contain SCNAs, several outstanding questions remain. For example, is gene compensation after SCNAs similar across tissue types? How do biological pathways, cellular localization and gene evolution influence the mechanism of gene regulation? Can we use SCNA analysis to investigate not only protein-level regulation but also RNA-level regulation and the relationship between the two? Here we perform a proteogenomic (DNA, RNA and proteins) analysis across primary tumor samples and cancer cell lines from different tumor types, a panel of isogenic non-tumorigenic human colon epithelial cells (hCEC) and normal tissues. We find tissue specificity in the RNA-level and protein-level compensation of genes affected by SCNAs. Importantly, we then utilize the DNA-RNA and RNA-protein correlations to infer the degree of regulation at the RNA and protein levels, respectively. In fact, as RNA-protein correlation informs us on the protein-level regulation, DNA-RNA correlation can inform us on the RNA-level regulation assuming DNA alterations are equally variable (as discussed below, see Figure 2B). Protein complex genes have a stronger protein-level regulation, while non-complex genes show stronger RNA-level regulation. Strikingly, we found an inverse relationship between the degree of RNA-level regulation and the degree of protein-level regulation across genes and cellular pathways. This suggests that cellular function impacts gene regulation and, for several pathways, tend to favor either RNA- or protein-level regulation. Finally, genes involved in RNA processing, translation and mitochondrial regulation are upregulated in highly aneuploid primary tumor samples (compared to low aneuploid tumors), especially at the protein level.

## Results

### Gene compensation at both the RNA and protein levels is specific to tissue type across tumors

Dosage compensation is a process by which cells modulate gene expression to buffer against changes in DNA copy number. In order to assess the degree of compensation at the RNA or protein level after DNA gains or losses (**Figure 1A**), we utilized the CPTAC dataset, a compendium of thousands of tumor samples analyzed for their genomic, transcriptomic and proteomic features (Ang et al., 2019). Here we analyzed most of the available CPTAC data comprising about 700 tumor samples derived from seven tumor types: colon adenocarcinoma (COAD), breast cancer (BRCA), ovarian cancer (OV), clear cell renal cell carcinoma (ccRCC), uterine corpus endometrial carcinoma (UCEC), head and neck squamous cell carcinoma (HNSC) and lung adenocarcinoma (LUAD). The dataset contains information at the DNA level through whole genome sequencing (WGS) or whole exome sequencing (WES), RNA level (RNAseq) and protein level (TMT Mass Spec) for 7-12K genes.

**Figure 1.**
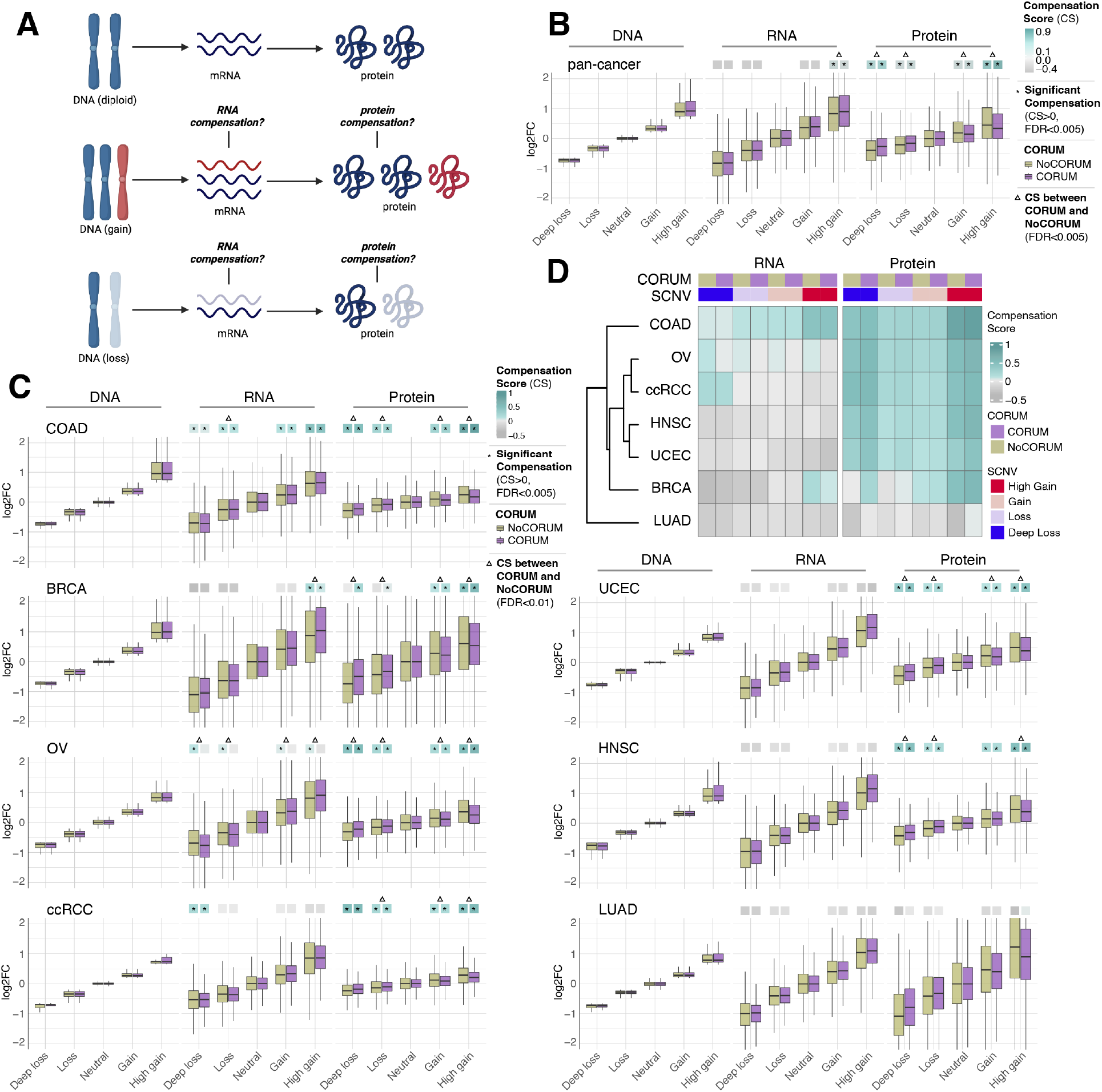
RNA-level and protein-level gene compensation across tumor types. (**A**)Schematic of RNA-level and protein-level gene compensation as a result of DNA gains (red) or losses (light blue). Dosage compensation is a process by which cells modulate gene expression to buffer against changes in DNA copy number. RNA and protein abundance change proportionally to the DNA change when dosage compensation is missing. (**B**) Box plots showing the CPTAC pan-cancer profile of DNA, RNA and protein log2FC in five groups based on the copy number change: deep loss (DNA log2FC < -0.65); loss (−0.65 < DNA log2FC < -0.2); neutral (−0.2 < DNA log2FC < 0.2), gain (0.2 < DNA log2FC < 0.65) and high gain (DNA log2FC > 0.65). The genes of each group were separated in protein complex genes (CORUM, purple) and non-complex genes (NoCORUM, yellow). The median of the compensation scores (CS) in each condition, which represents the degree of gene compensation, is shown at the top of the box plot (cyan/gray square). CS is positive when compensation happens (cyan) and is proportional to the degree of compensation. To test whether CS were significantly positive, we used bootstrapping test and p-values were corrected for FDR. An asterisk in the square indicates significant CS (CS>0 and FDR<0.005). A triangle above the squares indicates that the CS of complex and non-complex genes are significantly different by bootstrapping test (FDR<0.005). (**C**) Box plots showing the profiles of DNA, RNA and protein log2FC of the indicated cancer types grouped in five groups based on the copy number change as in **B**. The median CS is shown at the top of the box plots (cyan/gray squares). An asterisk in the square represented significant compensation (CS>0 and FDR<0.005). A triangle above the squares indicates that the CS of complex and non-complex genes are significantly different by bootstrapping test (FDR<0.005). (**D**) Heatmap showing the RNA-level and protein-level CS of different cancers. Cancers were clustered by Euclidean distance and hierarchical clustering.

For each gene of each cancer type, we defined the samples that did not have DNA copy number changes (log2 copy number ratio between -0.2 to 0.2) as the neutral group. We considered the median of the RNA and protein amount of this neutral group as the *neutral* RNA or protein level. Then, we calculated the log2 fold change (Log2FC) of the DNA, RNA and protein amount for each gene in each tumor sample relative to the corresponding *neutral* levels. We next determined the distributions of DNA, RNA and protein Log2FC of all genes from the 7 tumor types, based on 5 groups of DNA change (i.e., deep loss, loss, neutral, gain, high gain). Within each of these 5 groups, we also split by protein complex genes and non-complex genes based on the CORUM database (Ruepp et al., 2008) (see below, **Figure 1B**). In order to quantify the degree of RNA- or protein-level compensation, we calculated a compensation score (CS), for each gene in each sample, determined as the difference between the RNA or protein Log2FC and the DNA Log2FC (see Methods). To assess whether there was significant compensation in each group of DNA change (CS was significantly larger than zero), we implemented a bootstrapping method by randomly sampling the CS of genes within each group.

Work done in model organisms and isogenic human cells, suggests that RNA levels change proportionally to the DNA levels in aneuploid cells (i.e., there is no RNA-level compensation) while the protein levels do not (i.e., there is protein-level compensation; (Oromendia et al., 2012; Stingele et al., 2012; Torres et al., 2010)). In our pan-cancer analysis (**Figure 1B**), this was generally the case, with a significant protein-level compensation (FDR<0.001) and no significant RNA-level compensation except in the high gain group (genes showing high gain; see also below) (FDR<0.001). Protein-level compensation was significant for both gains and losses although was stronger in the high gain group versus the deep loss group (FDR<0.001).

Despite these overall trends in the pan-cancer analysis, we observed profound tissue specificity when we conducted the same analysis for each tumor type (**Figure 1C-D**). While protein-level compensation was widespread, LUAD did not show protein-level compensation (FDR=1) and BRCA showed reduced protein-level compensation compared to other cancer types, especially for DNA losses. Surprisingly, we found significant RNA-level compensation in certain tumor types. COAD showed general RNA-level compensation both for gains and losses (FDR <0.001; degree of compensation was lower for deep loss than other SCNA groups). In addition, ccRCC showed RNA-level compensation for deep losses (FDR<0.001), BRCA exhibited RNA-level compensation for high gains (FDR<0.001). Interestingly, and OV showed RNA-level compensation for non-complex genes but not for complex genes in all SCNA groups (see below).

Next, for each SCNA group (loss, deep loss, gain, high gain) we compared the genes belonging to protein complexes (“CORUM”, 3449 protein complex genes) with those that do not (“NoCORUM”, non-complex genes, i.e. remaining genes) using the CORUM dataset ((Ruepp et al., 2008); **Figure 1B-D**). We found that in general protein complex genes had a stronger protein-level compensation compared to non-complex genes (**Figure 1B**, FDR<0.001), consistent with previous studies examining the effect of chromosome gains (Oromendia et al., 2012; Stingele et al., 2012; Torres et al., 2010). Importantly, this was true not only for gains, but also for losses (**Figure 1B**, FDR<0.001) and across tumor types (**Figure 1C**, FDR<0.001). Interestingly, at the RNA level, the opposite was true. In the pan-cancer analysis protein complex genes showed less RNA-level compensation compared to non-complex genes for high DNA gain, the only group showing significant compensation in the pan-cancer analysis (**Figure 1B**, FDR<0.001). As mentioned above, for the individual tumor types, only certain cancers showed significant RNA-level compensation; in the majority of those cases complex genes showed less RNA-level compensation compared to non-complex genes (**Figure 1C**, FDR<0.001). For example, OV showed significant RNA-level compensation for non-complex genes but not for complex genes across all DNA groups (**Figure 1C**, FDR<0.001). In other words, protein complex genes showed changes at the RNA level that were more similar (in amplitude) to the changes observed at the DNA level, than non-complex genes, implying a lower level of regulation at the RNA level (see next section).

To validate the findings observed from primary tumors, we performed the same analysis on cancer cell lines using the Cancer Cell Line Encyclopedia (CCLE, (Barretina et al., 2012; Nusinow et al., 2020)), which showed general protein-level compensation and negligible RNA-level compensation except in deep loss group at the pan-cancer level (FDR<0.001). Protein complex genes have stronger protein-level compensation than non-complex genes similar to primary tumors (FDR<0.001). To further test this observation, we generated a panel of isogenic immortalized non-transformed hCEC with different aneuploidy patterns. We treated hTERT-immortalized TP53-KO (non-tumorigenic) hCEC (Martin et al., 2017) with reversine, an MPS1 inhibitor that inhibits correct chromosome attachment and spindle checkpoint, to induce random chromosome missegregation and subsequent aneuploidy (Santaguida et al., 2015). The single-cell derived clones contained different patterns of aneuploidy, characterized by WGS. We analyzed their transcriptome and proteome using RNA-sequencing and tandem mass spectrometry (MS/MS), respectively (see Methods). Interestingly, in addition to the widespread protein-level compensation, hCEC also showed RNA-level compensation as COAD did (FDR<0.001). Similar to COAD primary tumors, complex genes have stronger protein-level compensation (for DNA gain and deep loss group) but weaker RNA-level compensation (for DNA loss group) (FDR=0.013).

Overall, these data suggest that while protein-level compensation is widespread and RNA-level compensation is virtually absent in pan-cancer analysis, there is profound tissue specificity especially in the presence and degree of RNA-level compensation. Indeed, some tissue types (such as lung cancer) show low level of compensation both at the RNA and protein level, while others (as colon, breast, ovarian and renal cancer) show unexpectedly high compensation at the RNA level. Furthermore, protein complex genes generally showed stronger protein-level compensation and weaker RNA-level compensation compared to non-complex genes.

### Protein complex genes have a higher protein-level regulation and a lower RNA-level regulation than non-complex genes

Our previous analysis (**Figure 1B-D**) suggested that protein complex genes have stronger protein-level regulation and weaker RNA-level regulation compared to non-complex genes. To better understand this phenomenon, we decided to systematically study the correlation between DNA and RNA levels and between RNA and protein levels for each gene across samples, to infer the degree of gene regulation at the RNA and protein levels respectively (**Figure 2A**). In other words, if the correlation between RNA and protein is very high, we can infer that the protein abundance is mainly determined by the RNA amount with minimal protein-level regulation. On the other hand, if the correlation between RNA and protein is low, we assume a strong level of protein-level regulation. A similar logic can be used to infer the RNA-level regulation based on the DNA-RNA correlation, assuming DNA alterations are equally variable. To do this, we calculated Spearman’s correlation coefficients (rho) between DNA and RNA levels and between RNA and protein levels for each gene across tumor samples (**Figure 2B-2D**). Since the correlation coefficient depends on the extent of variation, we excluded the genes that show little or no changes at the DNA level across samples (−0.02< DNA log2FC <0.02 in more than 70% of the samples; see **Methods**). In addition, we also performed additional control analyses by analyzing the variance of DNA alterations (see below). We analyzed the pan-cancer distributions of the correlation coefficients for complex and non-complex genes using as the pan-cancer correlation the mean correlation coefficient value across the 7 tumor types (**Figure 2C**). As expected, we found that the median of the RNA-protein correlations was significantly lower for protein complex genes than for non-complex genes (FDR<0.001, bootstrapping test), indicating that protein complex genes tend to have stronger protein-level regulation compared to non-complex genes(Stingele et al., 2012). Strikingly, the opposite was true for RNA-level regulation, where the median of the DNA-RNA correlations was significantly higher for protein complex genes than for non-complex genes (FDR<0.001, bootstrapping test). This was in agreement with the observations described above regarding compensation in complex and non-complex genes (**Figure 1B-D**) and was not due to difference in the RNA abundance (analysis repeated with the exclusion of genes of low RNA abundance) or difference in the variance across DNA alterations (analysis of the variance across DNA values) between protein complex and non-complex genes. This result indicates that protein complex genes are likely to have weaker RNA-level regulation compared to non-complex genes. As a lower RNA-protein correlation suggests a stronger protein-level regulation, a lower DNA-RNA correlation similarly suggests a stronger RNA-level regulation. This RNA-level regulation could happen during transcription or other processes that influence RNA abundance. We also note that in terms of absolute correlation values (Spearman’s correlation), protein complex genes have lower RNA-protein correlations compared to DNA-RNA correlation values (median DNA-RNA correlation: 0.44; median RNA-protein correlation: 0.36 for protein complex genes, pan-cancer analysis), while it is the opposite for non-complex genes (median DNA-RNA correlation: 0.31; median RNA-RNA-protein correlation: 0.42 for non-complex genes, pan-cancer analysis, **Figure 2C**).

**Figure 2.**
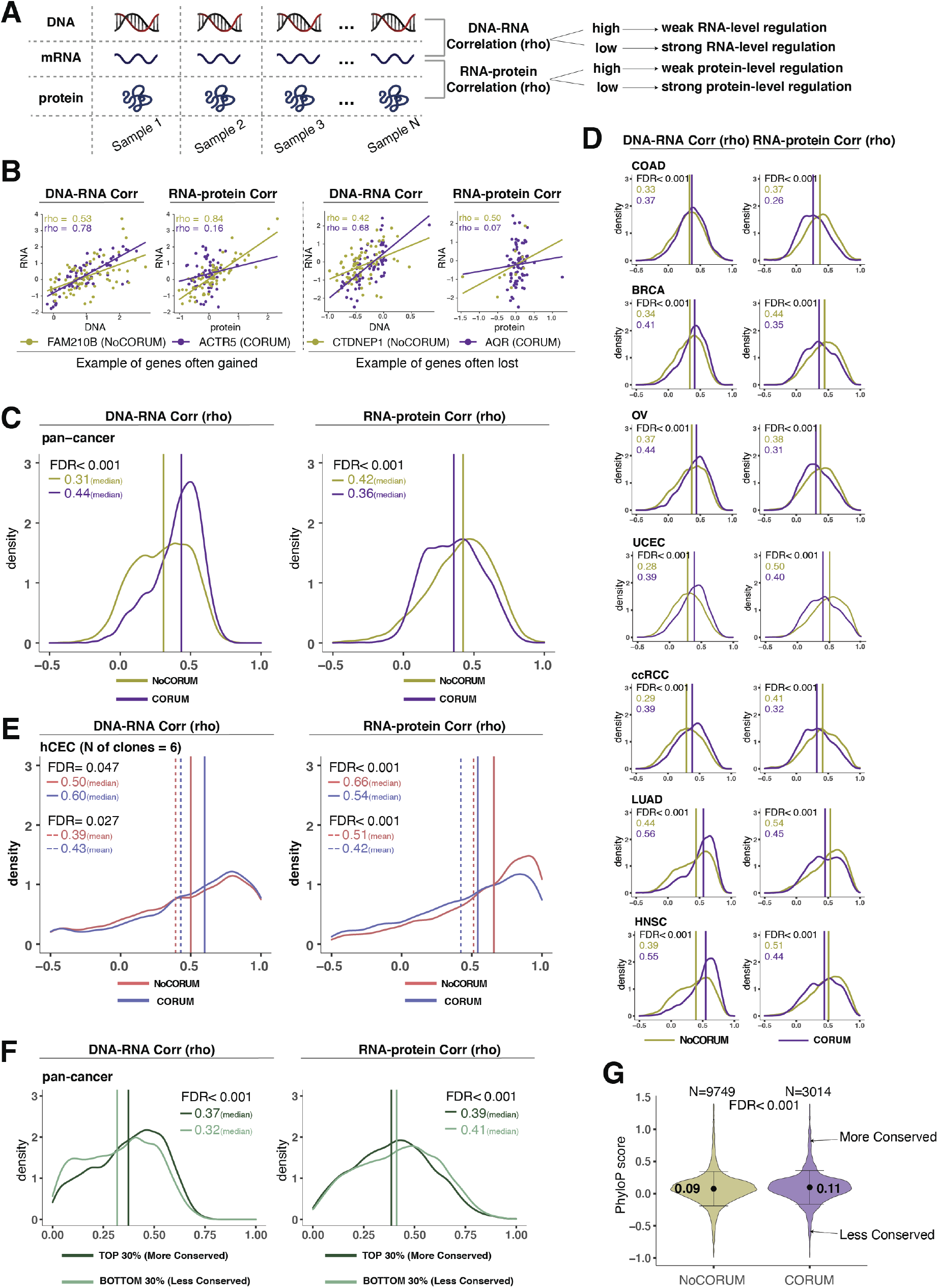
Protein complex genes have a stronger protein-level regulation than non-complex genes while non-complex genes have a stronger transcript-level regulation. (**A**)Schematic representing the strategy to infer the degree of RNA-or protein-level regulation by (Spearman’s) correlation analysis between DNA and RNA (DNA-RNA) or between RNA and protein (RNA-protein). A high (versus low) correlation is associated with a weak (versus strong) regulation. (**B**) DNA-RNA and RNA-protein correlations for representative CORUM and NoCORUM genes frequently gained (FAM210B and ACTR5) or lost (CTDNEP1 and AQR) in COAD. Dots represent different samples; solid lines indicate the linear regression line between DNA-RNA and RNA-protein; Spearman’s correlation is shown for each gene. (**C**) Density distribution of DNA-RNA and RNA-protein correlations for pan-cancer analysis (protein complex genes in purple and non-complex genes in golden yellow). Vertical lines and numbers in the top left represent the median correlation of protein complex genes (CORUM; purple) or non-complex genes (NoCORUM; golden yellow). Difference of the median correlation coefficients between protein complex genes and non-complex genes was evaluated by bootstrapping and p-values were adjusted for FDR. (**D**) Density distribution of DNA-RNA and RNA-protein correlations for individual CPTAC cancer types (protein complex genes in purple and non-complex genes in golden yellow). Purple vertical lines and numbers in the top left represent the median correlation of protein complex genes (CORUM). Golden yellow vertical lines and numbers in the top left represent the median correlation of non-complex genes (NoCORUM). Difference of the median correlation coefficients between protein complex and non-complex genes was evaluated by bootstrapping and p-values were adjusted for FDR. (**E**) Density distribution of DNA-RNA and RNA-protein correlations for hCEC cell lines (protein complex genes in blue and non-complex genes in red). Blue vertical solid (or dashed) lines and numbers in the top left represent the median (or mean) correlation of protein complex genes (CORUM). Red vertical solid (or dashed) lines and numbers in the top left represent the median (or mean) correlation of non-complex genes (NoCORUM). Difference of the median correlation coefficients between protein complex genes and non-complex genes was statistically evaluated by bootstrapping and p-values were adjusted for FDR. (**F**) Density distribution of DNA-RNA and RNA-protein correlations for evolutionally more conserved genes (dark green; genes in the top 30% of PhyloP scores) and less conserved genes (light green; genes in the bottom 30% of PhyloP scores). Dark green vertical lines and numbers in the top right represent the median of the more conserved genes; light green and numbers in the top right represent the median of the less conserved genes. Difference of the median correlation coefficients between more and less conserved genes was statistically evaluated by bootstrapping and p-values were adjusted for FDR. (**G**) The PhyloP score difference between protein complex genes (CORUM) and non-complex genes (NoCORUM). Difference between protein complex genes and non-complex genes was statistically evaluated by bootstrapping and p-values were adjusted for FDR.

We next extended these analyses to the 7 individual tumor types and we observed that this result was recapitulated across all of them (FDR<0.001, bootstrapping test, **Figure 2D**). Furthermore, this finding was confirmed using other proteogenomic datasets of cancer cell lines such as CCLE and NCI-60 (Alley et al., 1988). For the DNA-RNA correlation analysis, we also used the TCGA dataset (Cancer Genome Atlas Research Network et al., 2013) containing additional tumor types with DNA and RNA but without protein information and confirmed that DNA-RNA correlations were significantly higher for protein complex genes than for non-complex genes.

Next, we wanted to test whether these results obtained from primary tumors or cancer cell lines were recapitulated in non-tumor-derived cell lines and normal tissues. We tested whether this finding was recapitulated in our panel of isogenic untransformed hCEC with different aneuploid patterns. We confirmed that the DNA-RNA correlation was significantly higher and the RNA-protein correlation was significantly lower among protein complex genes than those among non-complex (**Figure. 2E**). Finally, we interrogated a database of normal tissues that includes RNA and protein levels for the RNA-protein correlations but not DNA copy number changes (as no SCNAs are generally present in normal tissues; (Wang et al., 2019)). Similarly, even in the normal tissues, we confirmed the lower RNA-protein correlation for complex genes compared to non-complex genes. Altogether these data indicate that protein complex genes have a stronger protein-level regulation, and a weaker RNA-level regulation compared to non-complex genes. These results also indicate that our findings in the tumors or tumor cell lines are recapitulated in untransformed isogenic aneuploid cells as well as normal tissues.

In addition to participation in protein complexes, we investigated other parameters, including biophysical properties and evolutionary conservation, for their association with gene regulation (DNA-RNA or RNA-protein correlation). Many of these properties, including protein regulatory sites, mRNA abundance, intrinsic protein disorder score and protein polarity, had no significant association with the type of gene regulation. The non-exponential degradation score, i.e. a score representing the likelihood that a protein is degraded in a non-exponential way (derived from (McShane et al., 2016)) was predictive of a strong regulation at the protein level, consistent with previous findings (McShane et al., 2016). Interestingly, the evolutionary conservation score (phyloP score) ((Hubisz et al., 2011); see Methods) was associated with the RNA-level and protein-level regulation. When we compared genes with high versus low conservation, we found that more conserved genes tended to have lower RNA-protein correlation and a higher DNA-RNA correlation compared to less conserved genes (**Figure 2F-G**; FDR<0.001). This indicates that more conserved genes tended to have a stronger protein-level regulation and a lower RNA-level regulation compared to less conserved genes. The conservation score (mean: 0.11 vs 0.09, FDR<0.001; variance: 0.31 vs 0.34, p=0.004) was also significantly higher for protein complex than non-complex genes (**Figure 2G**). Altogether, these data indicate that protein complex genes are more evolutionarily conserved and that the conservation score is associated with their preferred type of gene regulation (see Discussion).

### Negative association between RNA-level regulation and protein-level regulation across cellular pathways

We next performed a systematic analysis to understand whether and how gene function (i.e. belonging to a certain cellular pathway) and subcellular location influence the extent of RNA-level and protein-level regulation. This would inform us on whether or not genes may have evolved a preferred mechanism of gene regulation depending on the biological function or cellular distribution of the encoded protein. We started by examining the relationship between the DNA-RNA and the RNA-protein correlations across genes from the CPTAC tumors (pan-cancer analysis). Interestingly, we found that there was a significant negative association between these two parameters (slope=-0.33; association between *x* and *y* using the maximum of the density distribution of *y* along *x*: rho=-0.78, p=7.9E-07, **Figure 3A**, see **Methods**). In other words, genes showing a high DNA-RNA correlation tend to have a low RNA-protein correlation and vice versa. Based on this finding, we next asked whether the genes residing at the two ends of the distribution (high DNA-RNA correlation and low RNA-protein correlation, or low DNA-RNA correlation and high RNA-protein correlation) show enrichment in specific biological function. To this end, we defined two main groups of genes using the DNA-RNA and RNA-protein correlations (pan-cancer analysis): Group 1, composed of genes with a high DNA-RNA correlation (top 35%, rho>0.43) and a low RNA-protein correlation (bottom 35%, rho<0.31) and Group 2 of genes with a low DNA-RNA correlation (bottom 35%, rho<0.24) and a high RNA-protein correlation (top35%, rho>0.50). Gene Ontology (GO) enrichment analysis showed that Group 1 was strongly enriched in mitochondrial pathways (e.g. mitochondrial translation in pan-cancer; FDR=4.5E-43), protein translation (e.g. ribosome biogenesis; FDR=3.3E-23 and cytoplasmic translation in pan-cancer; FDR=1.0E-07) and RNA processing (e.g. RNA splicing; FDR=4.6E-29 and non-coding-RNA metabolism; FDR=6.0E-23). On the other hand, Group 2 was enriched in cell structure (e.g. actin filament organization; FDR=9.4E-10), cell adhesion (e.g. substrate adhesion, FDR =2.1E-17 and matrix adhesion, FDR=2.2E-11) and cell migration (FDR=0.044). Analysis in individual tumor types was overall similar to the pan-cancer analysis results for the pathways enriched in Group 1 and 2. Importantly, the result of the pathway enrichment analysis in the two groups of genes (Group 1 and 2) was validated using the CCLE dataset.

**Figure 3.**
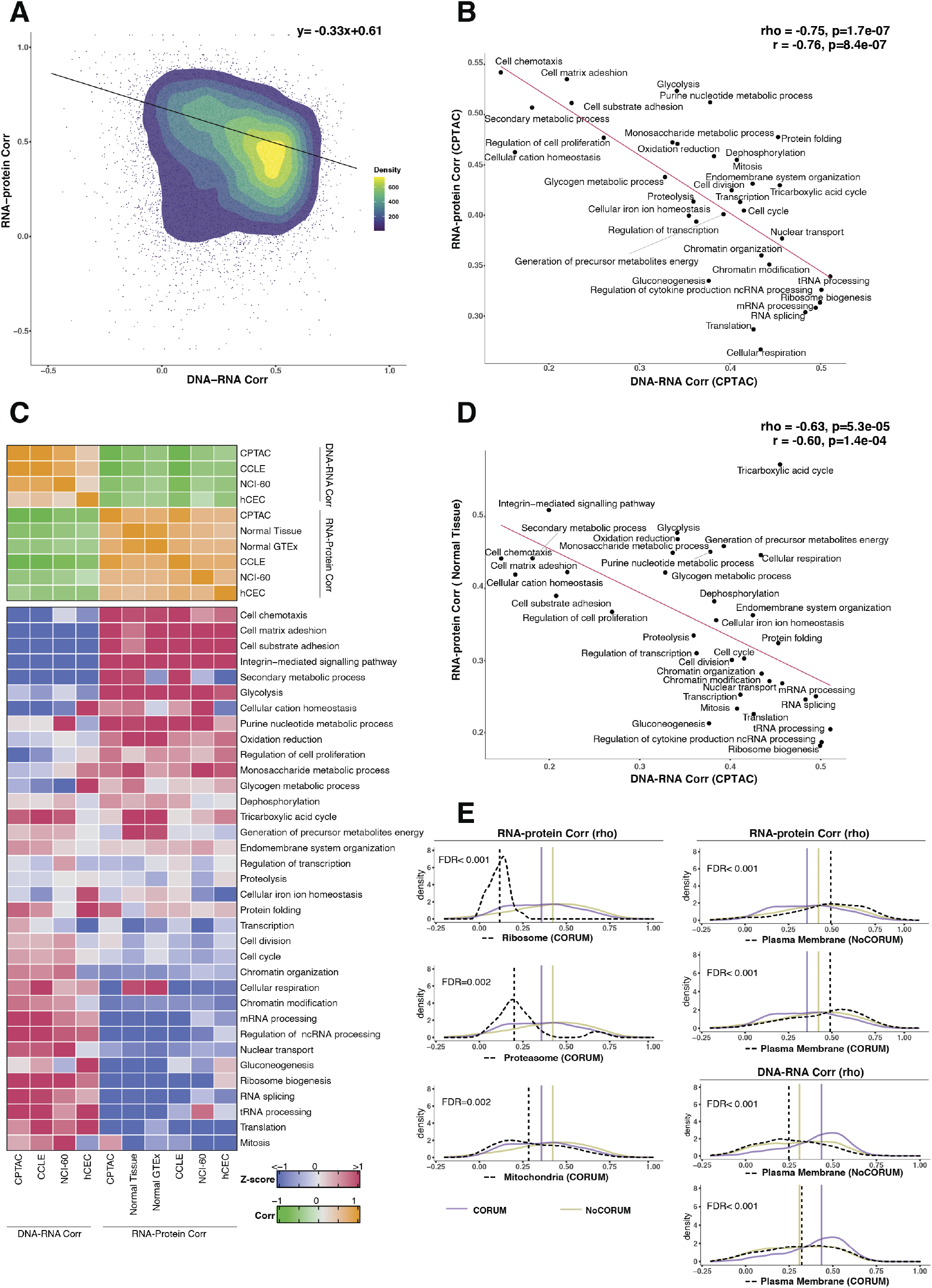
Functional characteristics of genes with different DNA-RNA and RNA-protein correlations: negative association between RNA- and protein-level regulation across pathways. (**A**) Dot plot where each point is a gene, DNA-RNA correlation is shown along the x-axis and RNA-protein correlation is shown along the y-axis. Density distribution is shown and a density distribution-dependent slope was calculated (see Methods) to estimate the association between the DNA-RNA and RNA-protein correlations. (**B**) A pathway-level analysis for the DNA-RNA and RNA-protein correlations (pan-cancer analysis, CPTAC). The DNA-RNA (x axis) and RNA-protein (y axis) correlation for each cellular pathway was calculated using the median rho value across all genes belonging to the pathway (pathway database: msigdbr, v7.4.1, category=C5). Spearman’s (rho) and Pearson’s (r) correlation coefficients are shown at the top right of the panel. (**C**) **Top panel**: a heatmap showing Spearman’s correlations among different proteogenomic datasets. For each dataset, we first calculated the DNA-RNA and RNA-protein rho values for each gene and then we calculated the Spearman’s correlation between these rho values (DNA-RNA rho or RNA-protein rho) across datasets. **Bottom panel**: a heatmap showing the pathway-level DNA-RNA and RNA-protein correlation score among different datasets. The pathway-level score was calculated by the median value across all genes in the same pathway and then Z-score transformed (pathway database: msigdbr, v7.4.1, category=C5). (**D**) A pathway-level analysis for the DNA-RNA (CPTAC) and RNA-protein (Normal tissues, Wang et al., 2019) correlations. The DNA-RNA (x axis) and RNA-protein (y axis) correlation for each cellular pathway was calculated using the median rho value across all genes belonging to the pathway (pathway database: msigdbr, v7.4.1, category=C5). Spearman’s (rho) and Pearson’s (r) correlation coefficients are shown at the top right of the panel. (**E**) Density distribution of DNA-RNA correlations for genes belonging plasma membrane and RNA-protein correlations for genes belonging to ribosome, proteasome, mitochondria and plasma membrane. The black dashed line represents the specific sublocation, the transparent purple line represents the median of CORUM complex genes and transparent golden yellow line represents the median of NoCORUM genes. Significance between the genes in the specific sublocation and all CORUM or NoCORUM genes was statistically evaluated based on bootstrapping test and adjusted for FDR. For example, the FDR in the top right panel was evaluated based on the difference between Ribosome (CORUM) and all genes in CORUM.

Since the genes showing similar DNA-RNA and RNA-protein correlations were enriched for specific cellular pathways suggesting similar biological function, we calculated the median value for these correlations among the genes in each pathway, thus obtaining pathway-level values for the DNA-RNA and RNA-protein correlations. For this analysis we considered the cellular pathways utilized in the Schwanhausser et al. 2011 (Schwanhäusser et al., 2011) using the msigdb GSEA database to identify the genes in each pathway (v7.4); altogether the genes in these pathways amounted for 84% of all genes. We found a strong negative correlation between the pathway-level DNA-RNA and RNA-protein correlations (rho=-0.75, p=1.7E-07; **Figure 3B**), corroborating and extending the finding described above at individual gene level. In agreement with our previous enrichment analysis, the pathway analysis showed RNA processing and translation pathways showed a preference to have high DNA-RNA and low RNA-protein correlations while cell adhesion and matrix-related pathways tended to have high RNA-protein and low DNA-RNA correlations (**Figure 3B-C**). Importantly, these results were confirmed based on the CCLE and NCI-60 datasets and based on our isogenic hCEC data (**Figure 3C**). Finally, we also confirmed the negative correlation trend was not determined by the variance across DNA values, i.e., by the fact that the genes in certain pathways were more likely to be gained or lost than the genes in other pathways. In addition, we also observed that RNA half-life was positively associated with the RNA-protein correlation (rho=0.51, p=0.001) and was negatively associated with the DNA-RNA correlation (rho=-0.52, p=0.002), while no association was found with protein half-life (see Discussion).

We next asked whether this result could also be recapitulated using proteogenomic datasets from normal tissues by interrogating two databases of normal tissues that include RNA and protein levels (GTEx Consortium, 2013; Jiang et al., 2020; Wang et al., 2019). By plotting the RNA-protein correlations calculated from normal tissues (instead of CPTAC) versus the DNA-RNA correlations from CPTAC, once again we found a significant negative correlation (rho=-0.63 p=5.3E-05; **Figure 3C-D**). This result indicates that there is an inverse relationship between the RNA-level and protein-level regulation across cellular pathways and that this is true also in normal tissues. Altogether these results suggest that RNA-level and protein-level regulation tend to be negatively associated across genes and pathways, perhaps due to an evolutionary selective pressure to favor one type of regulation over the other, depending on gene function and/or energetic demands of different cellular pathways (see Discussion).

Since specific biological pathways were predictive of whether a gene was more strongly regulated at the protein or RNA level, we wondered whether the subcellular localization of the gene was also related to the type of gene regulation. To test this, we split protein complex and non-complex genes into the following subcellular location/organelle groups: nucleus, nucleoli, cytoplasm, organelles, vesicles, Golgi apparatus, peroxisomes, lysosomes, endoplasmic reticulum, ribosome, proteasome, mitochondria and plasma membrane ((Thul et al., 2017), see **Methods**). We then used the DNA-RNA and RNA-protein correlations of genes in the same subcellular groups (calculated at the pan-cancer level) to determine whether the cellular localization was associated with different degree of RNA-level or protein-level regulation. We asked this question separately for complex and non-complex genes. Four subcellular locations/organelles stood out as different from the rest: ribosome, proteasome, mitochondria and plasma membrane (PM). In fact, among protein complex genes, those encoding for proteins taking part in the ribosome, proteasome and mitochondria showed a significantly lower RNA-protein correlation (suggesting stronger protein-level regulation) compared to protein complex genes in other cell locations (FDR<0.001 for ribosome; FDR=0.002 for proteasome and FDR=0.002 for mitochondria; **Figure 3E**). No difference was observed for non-complex genes of mitochondria compared to non-complex genes of other cell locations. Interestingly, the opposite behavior was observed for genes encoding for proteins located on the PM and this was true for both complex and non-complex genes. Protein complex or non-complex genes encoding for proteins located on the PM showed higher RNA-protein correlation and lower DNA-RNA correlation, compared to protein complex or non-complex genes encoding for proteins at other cell locations respectively, suggesting a significantly lower regulation at the protein level (FDR<0.001 for complex or non-complex genes; **Figure 3E**). This suggests that PM genes have a profoundly different type of regulation compared to other cellular locations, showing a low level of regulation at the protein level and a higher level of regulation at the RNA level (see Discussion). This is consistent with the GO enrichment analysis shown previously where cell structure and cell adhesion pathways were enriched in Group 2. Finally, GO enrichment analysis within PM genes with a low protein-level regulation showed an enrichment of cell substrate adhesion (FDR=1.5E-20) and cell leading edge (FDR=8.1E-34) including *ACTN1, CTNND1, DAG1* and others. Analysis within individual tumor types confirmed these results for the vast majority of cancer types.

### Protein-level changes associated with high levels of aneuploidy in primary tumors

While many studies have investigated the transcriptional changes associated with high level of aneuploidy in cancer, little is known about how these changes translate to the protein level, especially in primary tumors (Cancer Genome Atlas Research Network et al., 2013; Rodriguez et al., 2021). Given our finding on gene regulation across genes and pathways (**Figure 3**), it is likely that the dysregulation of certain pathways in cancer may be overlooked by investigating exclusively changes at RNA level, as most studies have done. Thus, we set out to investigate which pathways are enriched (increased expression) or depleted (decreased expression) in highly aneuploid tumors compared to tumors with low aneuploidy both at the protein level and at the RNA level (**Figure 4A**). We note here that the goal is to identify expression changes resulting from higher aneuploidy independent of the specific chromosomes that are gained or lost. We first determined the overall aneuploidy level (aneuploidy score) for each primary CPTAC tumor by calculating the total number of chromosome arms gained or lost (across all chromosomes) (see also Methods). Second, we used a linear regression model to study the association between the RNA or protein level of each gene and the aneuploidy score. Genes were ranked based on the t-value associated to the aneuploidy score and GSEA was performed on the ranked gene lists to assess which pathways were differentially expressed according to the aneuploid score (see also Methods). Finally, in order to take into account potential confounding variables (such as tumor purity and cell cycle score), we repeated the linear model including them as covariates. For the pan-cancer analysis, we also included the tumor type as a covariate.

**Figure 4.**
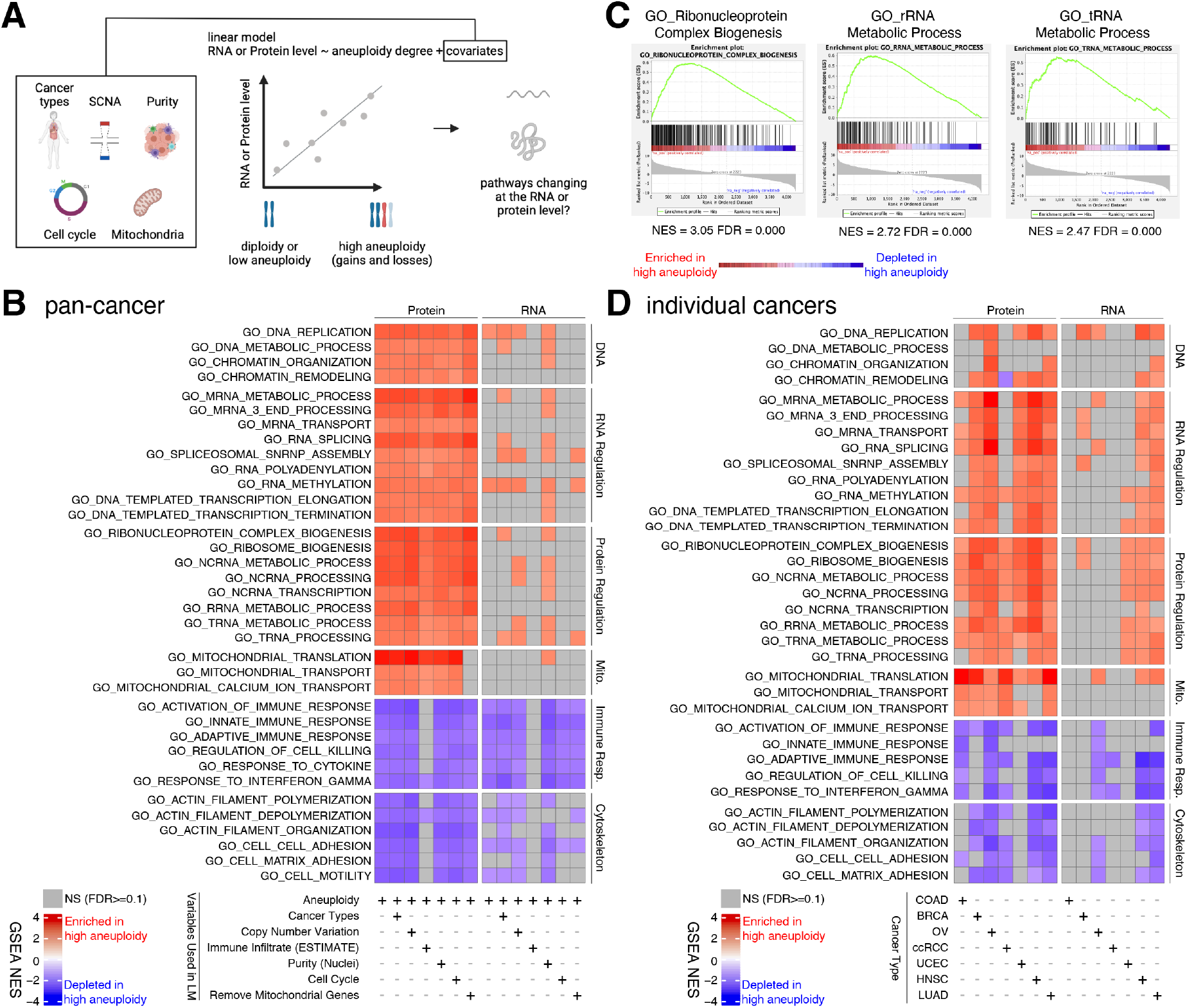
Analysis of pathways dysregulated at the RNA and protein levels in high aneuploidy tumor samples. (**A**) Schematic of the method used to identify pathways changing at the RNA and protein levels in samples of high aneuploidy. The aneuploidy degree of primary tumors (CPTAC) was used to fit the RNA or protein level of each gene by linear models. Several covariates were included in the model one by one to exclude their influence on the RNA or protein level, including cancer type, gene-level copy number variation, purity, cell cycle and mitochondria. T-values associated to the aneuploidy score was used to rank genes for gene set enrichment analysis (GSEA). (**B**) A heatmap showing the enrichment score for the indicated pathways significantly enriched (red) or depleted (blue) in high versus low aneuploidy tumor samples. Specific gene sets related to DNA, RNA and protein regulation and mitochondria are enriched and those related to immune response and cytoskeleton are depleted at the protein level in aneuploid tumor tissues. Covariates were included in the model to control for cancer types, gene-level copy number variation, purity and cell cycle scores. Mitochondrial genes were removed in the last column. The gene sets whose FDR are larger than 0.1 are shown in gray. (**C**) Enrichment plots of three pathways related to protein translation in tumor tissues: ribonucleoprotein complex biogenesis, rRNA metabolic process and tRNA metabolic process. The normalized enrichment scores and FDR are shown below the corresponding enrichment plots. (**D**) A heatmap showing the enrichment of the same gene sets as (B) in individual cancer types. RNA or protein expression for each gene was fit by the aneuploidy degree without the inclusion of other covariates. Gene sets enriched in high aneuploid samples are in red while those depleted in high aneuploid samples are in blue. The gene sets whose FDR are larger than 0.1 are shown in gray.

Pan-cancer analysis (variables in the model: aneuploidy score + cancer type) revealed several pathways to be enriched at the protein level in highly aneuploid tumors (**Figure 4B**). They included pathways related to DNA and chromatin such as DNA replication (GO: DNA replication, NES=2.78, FDR<0.001) and chromatin organization (GO: Chromatin organization, NES=2.36, FDR<0.001), RNA production and processing such as transcription elongation (GO: DNA-templated transcription elongation, NES=2.35, FDR<0.001), termination (GO: DNA-templated transcription termination, NES=2.37, FDR<0.001), RNA splicing (GO: RNA splicing, NES=2.93, FDR<0.001), RNA polyadenylation (GO: RNA polyadenylation, NES=2.33, FDR<0.001) and RNA transport (GO: mRNA transport, NES=2.35, FDR<0.001), pathways related to protein translation including rRNA (GO: rRNA metabolic process, NES=2.72, FDR<0.001), tRNA processing (GO: tRNA metabolic process, NES=2.47, FDR<0.001) and ribosome biogenesis (GO: Ribosome biogenesis, NES=2.85, FDR<0.001), and pathways related to mitochondrial gene expression (GO: Mitochondrial translation, NES=3.49, FDR<0.001) and transport (GO: Mitochondrial transport, NES=2.35, FDR<0.001) (**Figure 4B-C**). On the other hand, several pathways were also depleted in highly aneuploid tumors such as cytoskeleton (GO: Actin filament organization, NES=-2.54, FDR<0.001) cell adhesion (GO: Cell matrix adhesion, NES=-2.30, FDR=1E-04), and pathways related to immune responses (GO: Activation of immune response, NES=-2.57, FDR<0.001) (**Figure 4B**), consistent with previous studies (Davoli et al., 2017). Importantly, these results were confirmed after including additional covariates in the model (**Figure 4B**). First, we included tumor purity, which was estimated using two independent methods (nuclei percentage or using the algorithm Estimate;(Yoshihara et al., 2013)), confirming that the results were independent of the immune and stromal component of the tumor samples. Second, we included the DNA copy number change for each gene, in order to assess whether the change at the protein level associated with aneuploidy was due to the fact the genes change at the DNA level (gained or lost) or instead to transcriptional/translational programs that are established in high aneuploid cells. The result suggested that the latter is correct as it was confirmed after including the DNA copy number change information. Furthermore, as gene sets related to transcription and translation also include many mitochondrial ribosome, rRNA and tRNA genes, we removed all mitochondrial genes (mitochondrial genes encoded by the nuclear or mitochondrial DNA) before GSEA to exclude the possibility that these transcription and translation pathways were enriched only because of mitochondrial genes. The result of this analysis validated once again the fact that transcription and translation of nuclear DNA-encoded genes are upregulated in high aneuploid cancers. Finally, since our analysis found the cell cycle pathway as one of the enriched pathways at the protein level in high aneuploidy tumors (GO: Cell cycle checkpoint, NES=1.83, FDR=0.01), consistent with previous findings (Carter et al., 2006; Davoli et al., 2017), we repeated the linear model including the cell cycle score ((Davoli et al., 2017); see Methods) to assess changes in pathways independently of cell cycle change. This model suggested that the results observed in the original model were independent of cell cycle score. Analyses of individual tumor types were generally consistent with these results (**Figure 4D**).

Interestingly, when we repeated the same analyses using RNA level using the same set of genes, we observed a similar trend where pathways related to DNA, RNA and protein regulation and mitochondrial translation and transport were enriched while those related to immune response and cytoskeleton were depleted in tumor tissues of high aneuploidy (**Figure 4B and 4D**). However, the enrichment of pathways related to DNA, RNA and protein regulation and mitochondrial translation and transport at RNA level in tumor tissues was not as strong as the ones observed at protein level (more pathways were not significant at the RNA level compared to the protein level in **Figure 4B and 4D**). This difference between transcriptome and proteome is consistent with our findings in **Figure 3C** that pathways related to transcription, translation and mitochondria show a preference to be regulated at protein level.

Altogether these results suggest that tumors with high degree of aneuploidy show enrichment in pathways related to protein translation, mitochondria and RNA processing, which is independent of other covariates such as purity and cell cycle score. These changes are more evident at the protein level than at the RNA level suggesting that their upregulation is due at least in part to a protein-level regulation.

## Discussion

How cells control the abundance of their proteins in physiological and pathological conditions is a fundamental question. Both the regulation at the RNA and protein level can contribute to the protein abundances. However, the relative contribution of these two layers of regulation remains unclear (Vogel & Marcotte, 2012). Furthermore, it remains unknown whether the relative contribution of the RNA- and protein-level regulation vary based on the DNA copy number, interaction with other proteins, protein function and location and so on. In this study, our proteogenomic analysis allowed us to uncover general principles linking gene function and other features to the mechanism of gene regulation. In particular, we found that the genes and pathway that have a stronger protein-level regulation tend to have a weaker RNA-level regulation and vice versa, suggesting that each pathway has a predominant type of regulation.

### Tissue specificity of RNA- or protein-level compensation

Pan-cancer analysis revealed several forms of gene compensation that are common across the majority of tissue types (**Figure 1B**). We found that strong compensation at the protein level is generally widespread, while compensation at the RNA level is rare. The protein-level compensation is stronger for genes in protein complexes than non-complex genes, consistent with previous reports (Stingele et al., 2012; Torres et al., 2010). Interestingly, the existence of protein-level compensation and its higher degree for protein complex genes were true not only for DNA gains, but also for DNA losses. Consistent with our findings, a recent study reported protein-level compensation after chromosome loss (Chunduri et al., 2021) although in this study no significant difference was reported between complex and non-complex genes, perhaps due to the limited number of genes on the lost chromosomes. The protein compensation for complex genes of DNA gains is thought to occur through protein degradation of the overabundant subunits (McShane et al., 2016). However, this model cannot easily explain how protein compensation happens after DNA losses and why the compensation is stronger for protein complex genes. Future studies are needed to shed light on this process.

Individual tumor types showed unexpected tissue specificities for type and degree of compensation (**Figure 1C**). For example, lung adenocarcinoma did not show compensation, neither at the protein nor at the RNA level. Renal cancer and breast cancer showed RNA-level compensation for deep losses and for high gains, respectively. Furthermore, colon adenocarcinoma showed RNA-level compensation both for gains and losses. Ovarian cancer showed a similar general compensation at RNA-level but only for non-complex genes and not for complex genes (see also below). To our knowledge, this is the first study to investigate and report tissue-specific RNA- and protein-level compensation across different tumor types.

### Negative association between protein-level and RNA-level regulation across genes and pathways: their regulation tends to occur either at the RNA or protein level

In this study, we use the DNA-RNA correlation and the RNA-protein correlation as a way to estimate the degree of RNA-level and protein-level regulation. We observed that genes with similar pattern of regulation tended to be enriched in functional pathways thus to perform related functions. For example, genes implicated in translation and RNA processing tended to have stronger protein-level regulation while genes functioning in cell structure and adhesion tended to have lower protein-level regulation (Group 1 and 2 analysis). The opposite was true for RNA-level regulation, which was stronger in genes functioning in cell structure and adhesion. This indicates that genes sharing similar biological functions may have evolved similar types of regulation. Interestingly, a previous study investigating the RNA and protein half-lives reported functional similarities among pathways with similar RNA and/or protein half-lives (Schwanhäusser et al., 2011).

Strikingly we observed a significant negative correlation between the RNA-level and the protein-level regulation across both genes (**Figure 3A**) and cellular pathways (**Figure 3B**). This finding held true if we used normal tissue datasets to calculate the RNA-protein correlations (**Figure 3C**). This suggests that the degree of RNA-level regulation tend to be inversely associated with the degree of protein-level regulation. *Why is this the case?* One possibility is that there is an energetic constraint in the evolution of gene regulation. Given the high energetic demand of both RNA and (especially) protein synthesis (Wagner, 2005), it may be energetically favorable for the cells to have one predominant level of gene regulation for most pathways, at the mRNA or protein level. For example, genes functioning in protein complexes require a sophisticated and energy demanding mechanism of co-translational assembly to build a complex with the right subunit stoichiometry (Kamenova et al., 2019; Shiber et al., 2018; Taggart et al., 2020; Taggart & Li, 2018). This level of regulation would likely be still needed even if all the genes were transcribed in the right proportion. Thus it might be more beneficial for the cell to invest more energy on protein-level regulation and save energy at RNA-level one. On the other hand, for proteins involved in cell adhesion, often secreted or at the plasma membrane, it may be more difficult if not impossible (for the secreted ones) to regulate their level (for example through degradation) once synthetized and transported to the location where they normally function. Thus in this case, a strong RNA-level regulation may be necessary. Another possibility is the different demands of dynamic regulation for different pathways. For example, pathways controlling cell adhesion and migration may have a stronger need to rapidly regulate their function compared to translation or splicing pathways. Since RNA synthesis alters the abundance of transcripts at a higher rate than protein synthesis alters protein abundance (McManus et al., 2015), stronger RNA-level regulation (compared to protein-level) may be necessary for a timely and efficient protein production. A final possibility is the cellular localization of the proteins. For example, genes encoding mitochondrial proteins have a strong protein-level regulation. As proteins are synthetized before import into mitochondria (Isaac et al., 2018), regulation of protein function and complex assembly has to occur at the protein level within the organelle.

### Types of gene regulation and other gene features: cellular localization and mRNA half-life

Protein localization, which relates to protein function was also a predictor of the type of gene regulation. Ribosome and proteasome complexes showed the strongest level of protein-level regulation, consistent with previous observations (Taggart et al., 2020). Mitochondrial genes belonging to protein complexes showed a similarly strong protein-level regulation. On the contrary and very surprisingly, compared to the other cell locations, proteins that reside on the plasma membrane (PM) showed a weaker protein-level regulation and a stronger RNA-level regulation. While we cannot exclude that this may be due to technical difficulties in detecting membrane proteins, if this was the case, we would expect the mRNA-protein correlation to be lower (due to higher detection noise) and not higher than the other complex genes, as we instead observed (**Figure 3E**). While misfolding-induced degradation of proteins in the cytosol or ER (e.g. through the unfolded protein response) is well understood, very little is known about the consequence of misfolding or mis-assembly for proteins on the plasma membrane (Hetz et al., 2020).

We noticed an interesting association also with RNA half-life. RNA half-life was positively associated with the RNA-protein correlation (rho=0.508, p=0.001) and negatively associated with the DNA-RNA correlation (rho=-0.516, p=0.002). In other words, pathways with a strong protein-level regulation (Group 1, **Figure 3A**) tended to have a low RNA half-life and pathways with a strong RNA-level regulation (Group 2, **Figure 3A**) tended to have a high RNA half-life. Since pathways that tended to be strongly regulated at the RNA level have long lived RNA, this suggests that most regulation is at the transcriptional, not the RNA degradation level. The association with RNA half-life may be also related to the differential energetic demand of different pathways (Wagner, 2005; see above). Genes that have a strong RNA-level regulation (e.g. cell adhesion and migration) are perhaps more likely to invest significant energy in the fine-tuning of transcripts’ abundance; thus it may be beneficial for these transcripts to have longer half-lives given the energy consumption at the RNA-level regulation compared to transcripts derived from genes mainly regulated at the protein level.

### Pathways dysregulated in aneuploid cancers at the protein level

We observed that among the pathways significantly upregulated in high versus low aneuploid tumors (both pan-cancer and individual tumor type analyses), there were pathways related to RNA transcription, processing, transport and regulation, tRNA and ribosome biogenesis, and protein synthesis and translation. These gene sets are enriched among those that tend to have stronger protein-level regulation. Interestingly, in the flagship endometrial CPTAC study, ribosome biogenesis was one of the most strongly enriched pathways in the serous uterine cancer subtype (Dou et al., 2020), which is the one that shows the highest level of aneuploidy among all uterine cancer subtypes. We also note that, based on recent studies, most of these pathways that we found upregulated in primary tumors were not significantly enriched in high aneuploid cancer cell lines, based on recent reports (Schukken & Sheltzer, 2021). This suggests that the tumor microenvironment may play an important role in shaping the level of these pathways.

### Open questions

An outstanding question remains about what is the mechanism of protein-level compensation and regulation. Previous studies suggest that the regulation occurs at the level of protein degradation (Dephoure et al., 2014; Torres et al., 2010). However it seems now clear that protein degradation co-exists with regulation at the protein synthesis level and that at least for certain complexes, the vast majority of the protein level regulation occurs at the protein synthesis level with fine tuning happening through protein degradation (Kamenova et al., 2019; Shiber et al., 2018; Taggart et al., 2020; Taggart & Li, 2018). Additional studies are needed to better characterize the level of translation or proteasome regulation across cell location protein complexes and cellular pathways.

## Methods Datasets

All CPTAC-related SCNA, mRNA, protein and mutation data were from website (https://cptac-data-portal.georgetown.edu/datasets or directly from Clinical Proteomic Tumor Analysis Consortium) or directly from Clinical Proteomic Tumor Analysis Consortium.

The list of genes in protein complexes was from the CORUM database v3 (Core complexes) from http://mips.helmholtz-muenchen.de/corum/#download (Ruepp et al., 2008).

All CCLE-related SCNA, gene-expression and protein data for cell lines were derived from DepMap (CCLE_segmented_cn.csv, DepMap Public 19Q4; CCLE_gene_cn.csv, DepMap Public 19Q4 and CCLE_expression_full.csv, DepMap Public 19Q4; protein_quant_current_normalized.csv, DepMap Public 19Q4), respectively.

NCI-60-related data was from NCI (https://discover.nci.nih.gov/cellminer/loadDownload.do; DNA: Combined aCGH; RNA: RNA-seq; Protein: SWATH (Mass spectrometry)), respectively.

The 29 human tissue samples used for mRNA and protein expression analysis was from (Wang et al., 2019)

GTEx RNA data was downloaded from GTEx portal for RNA values (v8, https://gtexportal.org/home/), and for protein data was downloaded from (Jiang et al., 2020)

### Calculation of log2FC for DNA, RNA and protein values

Before starting the calculation, low-expression genes whose RNA level were within the bottom 10% in individual tumor tissues were removed. Only genes which had DNA, RNA and protein data were kept for the following analyses. For each gene of each cancer (80-110 patients for each cancer), we defined the patients that do not have a DNA copy number change (log2 copy number ratio is between -0.2 to 0.2) as the neutral group. We considered the RNA and protein expression median of this group as the *neutral* RNA or protein level. Then we calculated the log2 fold change (Log2FC) at the DNA, RNA and protein level for each gene in each sample compared to the *neutral* DNA, RNA or protein level. For each gene in each sample, we determined whether there is a DNA loss (DNA log2FC is between -0.65 to -0.2), deep loss (DNA log2FC < - 0.65), gain (DNA log2FC is between 0.2 to 0.65) or high gain (DNA log2FC > 0.65). For the pan-cancer analysis, the log2FC data of individual cancers were pooled together (682 patients in total). Before pooling together data from different tumor types, quality control was done using principal component analysis on the pooled log2FC data, confirming that no cancer type was separate from others (in the principal component analysis). To calculate the log2FC of cancer cell lines (CCLE), the cancer types of more than 13 cell lines were used (284 samples from 11 cancer types). As for CPTAC, the log2FC of DNA, RNA and protein were calculated for each gene in each cancer. Then the log2FC values of different cancers were merged.

### Calculation of compensation score

In order to quantify the degree of RNA- or protein-level compensation, we calculated a compensation score (CS) for each gene in each sample determined as the difference between the RNA or protein log2FC and the DNA log2FC as shown in the following formula. CS is larger than 0 when compensation exists. A higher CS means higher compensation.

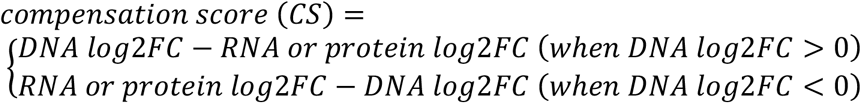

To test whether there was significant compensation in each group of DNA change, we used bootstrapping method by randomly sampling the CS of genes in the specific groups for 10000 times and calculated the median of CS for each time by boot package (v1.3-28). 95% confidential interval of CS was calculated by basic method of boot.ci function. The p-value was calculated at one-tail to test the H0: the CS is not larger than 0, which was corrected by FDR method. To compare whether there was significant difference between CS of protein complex genes and non-complex genes in the specific groups, the CS was randomly resampled for 10000 times and the difference of CS was calculated for each time by boot package. 95% confidential interval of CS difference (CS for protein complex genes – CS for non-complex genes) was calculated by basic method of boot.ci function (positive values mean stronger compensation for protein complex genes). The p-value was calculated at two-tail to test the H0: the CS difference equals 0, which was corrected for FDR Benjamin-Hochberg method.

### DNA-RNA and RNA-protein correlation for each gene

Only the genes which had DNA, RNA and protein data were considered for these analyses. For each cancer type, genes that showed no or very little change at the DNA level (−0.02< DNA log2FC <0.02) in more than 70% of the patients were removed because for those genes very little or no variance at the DNA level is likely to influence the correlations analyses. The analyses were also repeated using all genes and the results were confirmed. For each gene we then calculated the DNA-RNA and RNA-protein Spearman’s correlation (rho value). Next, we merged the correlation of different tumor types at the gene level. More specifically, for each gene, we calculated the mean of correlation coefficients across different tumors and considered this value as the correlation coefficient for the pan-cancer. The same method was applied to CCLE and NCI-60 datasets and to normal tissues datasets (Alley et al., 1988; Barretina et al., 2012), Then we calculated the RNA-protein Spearman’s correlation (rho value) for each gene. The p-value was evaluated based on a 10000-times bootstrapping test to compare the median difference between CORUM complex genes with NoCORUM genes. All the p-values were adjusted by FDR using the Benjamin-Hochberg method (Benjamini & Hochberg, 1995).

### Gene-level and pathway-level analysis of the DNA-RNA and RNA-protein correlations

We first calculated the pan-cancer DNA-RNA and RNA-protein Spearman’s correlations (rho values) for each gene as described above. In order to estimate the association between the DNA-RNA (DR) and RNA-protein (RP) correlations, for each DR find RP corresponding to the maximum of the density distribution. If the density distribution is f(DR, RP), we divide the DR range into n=40 bins, and for each DR_i_, we find

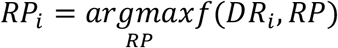

We then find the slope of (DR_i_,RP_i_), i=1..n, and use it as an estimate of the association between DNA-RNA and RNA-protein correlations.

For each cancer type, genes were divided into 2 groups: Group 1, composed of genes with a high DNA-RNA correlation (top 35%, rho>0.43) & a low RNA-protein correlation (bottom 35%, rho<0.31) and Group 2 of genes with a low DNA-RNA correlation (bottom 35%, rho<0.24) & a high RNA-protein correlation (top35%, rho>0.50). Gene Ontology enrichment analysis was then used to test whether these genes showed enrichment or not for different pathways (msigdbr, v7.4.1, category=C5).

For the pathway-level analysis, we first downloaded the gene sets for each pathway from GSEA (v7.4, https://www.gsea-msigdb.org/gsea/msigdb/collections.jsp#C3) and utilized the cellular pathways based on (Schwanhäusser et al., 2011). For each pathway, the median of the rho values across the genes in the pathway was utilized as the correlation value associated to the pathway (for example, the median of DNA-RNA rho correlation values for the genes in the Cell Cycle pathway would represent the DNA-RNA correlation value for Cell Cycle pathway).

### Phylogenetic conservation analysis

PhyloP scores (hg19.100way.phyloP100way.bw (Hubisz et al., 2011)) was downloaded from UCSC (http://hgdownload.cse.ucsc.edu/goldenpath/hg19/phyloP100way/, positive value: more conserved; negative value: less conserved)). Genome related information was downloaded from Genecode (http://ftp.ebi.ac.uk/pub/databases/gencode/Gencode_human/release_19/gencode.v19. annotation.gff3.gz). In order to transform the score from coordinates to gene-level, we first split the PhyloP scores from genome coordinates to gene coordinates and then calculated the median of PhyloP score along all the gene body as the PhyloP score for each gene. During these analyses, genes with top 30% and bottom 30% of the phyloP score were picked for the further analysis.

### Subcellular location analysis

Subcellular location data (subcellular_location.tsv) was downloaded from https://www.proteinatlas.org/about/download) (Uhlén et al., 2015). The “Main location” data was used for the Subcellular location analysis. Subcellular locations included: Nucleoplasm, Nuclear speckles, Nuclear bodies and Nuclear membrane were organized as Nucleus; Microtubules, Cytosol, Actin filaments, Centrosome, Centriolar satellite, Cytoplasmic bodies, Intermediate filaments, Cytokinetic bridge, Mitotic spindle and Microtubule ends were grouped as Cytoplasm; Nucleoli, Nucleoli fibrillar center and Nucleoli rim were the Nucleoli; Mitochondria, Endoplasmic reticulum, Plasma Membrane, Proteasome and Ribosome. For each subcellular location, we calculated the DNA-RNA and RNA-protein Spearman’s correlation for the genes in each subcellular location. The p-value was evaluated based on a 10000 times bootstrapping test. All the p-value was adjusted by FDR method.

### Generate single-cell derived hCEC clones containing aneuploidy

To derive a panel of isogenic aneuploid cell lines, hTERT-immortalized TP53-KO (non-tumorigenic) hCEC cells (Martin et al., 2017) were treated with reversine (0.2uM for 24 hours), an MPS1 inhibitor that prevents correct chromosome attachment and spindle checkpoint to induce random chromosome missegregation (Santaguida et al., 2015). Then the cells were plated at a low density and grew until the colonies formed. Those single-cell derived clones were picked using glass cylinders. To identify the levels and patterns of aneuploidy, the clones were sequenced by shallow whole genome sequencing. The transcriptome and proteome were measure by RNA sequencing and mass spectrometry (see below).

### Shallow whole genome sequencing

Low-pass (∼0.1-0.5X) whole-genome sequencing reads of hCEC were aligned to reference human genome hg38 by using BWA-mem (v0.7.17) (H. Li & Durbin, 2009) and followed by duplicate removal using GATK (Genome Analysis Toolkit, v4.1.7.0) (https://gatk.broadinstitute.org/hc/en-us) to generate analysis-ready BAM files. BAM files were processed by the R Package CopywriteR (v1.18.0) (Kuilman et al., 2015) to call the arm-level copy numbers.

### RNA sequencing

hCEC clones were plated in 6-well plates one day before the collection. At the second day, the cells were checked to make sure their confluency were within 70-90% and morphology was normal. Then the cells were wash twice by PBS and stored at -80C immediately. Total RNA was isolated from each sample using PicoPure RNA Isolation kit (Life Technologies, Frederick, MD) including the on-column RNase-free DNase I treatment (Qiagen, Hilden, Germany) following the manufacturers’ recommendations. To purify RNA for sequencing, we used the Qiagen RNeasy Mini Kit (Qiagen 74106). RNA concentration and integrity were assessed using a 2100 BioAnalyzer (Agilent, Santa Clara CA). Sequencing libraries were constructed using the TruSeq Stranded Total RNA Library Prep Gold mRNA (Illumina, San Diego CA) with an input of 250ng and 13 cycles final amplification. Final libraries were quantified using High Sensitivity D1000 ScreenTape on a 2200 TapeStation (Agilent, Santa Clara CA) and Qubit 1x dsDNA HS Assay Kit (Invitrogen, Waltham MA). Samples were pooled equimolar with sequencing performed on an Illumina NovaSeq6000 SP 100 Cycle Flow Cell v1.5 as Paired-end 50 reads.

### RNAseq pipeline

Total RNA sequencing reads of hCEC were mapped to the human genome hg38 by STAR (version 2.7.7a) (Dobin et al., 2013) using the 2-pass model. Hg38 sequence and RefSeq annotation were downloaded from the UCSC table browser. RSEM (version 1.3.1) (B. Li & Dewey, 2011) was used to quantify genes and transcripts expression levels. RSEM output the gene-level raw counts and FPKM (Fragments Per Kilobase of transcript per Million mapped reads) results in table format. The RNA RSEM data will be filtered for genes with median FPKM > 1 for use in downstream analyses.

### Global protein abundance profiling

Cell pellets were lysed in 8M urea 100 mM TRIS pH=8.5 + 10 mM TCEP + 40 mM CAA (150 ul/sample) by sonication in probe sonicator for 1 × 5 sec cycle @ amplitude of 50%. Lysates were incubated for 30 min @ 56oC in the thermoshaker @ 1,000 rpm. Insoluble debris were removed by centrifugation (5 min @ 16,000 x g). Protein concentrations were measured by A280 method and proteins were digested with trypsin @ 50:1 (w/w) ratio o/n @ 37oC (lysates were diluted 6 fold with 20 mM TRIS, pH=8 prior to digestion). Subsequently samples were acidified with 10% FA to final of 0.5% FA and centrifuged to remove undigested material. Peptides were desalted on tC18 Waters SepPak cartridges and eluates were dried on speedvac.

50 ug of digest from each sample were resolubilized in 20 ul of 50 mM HEPES buffer pH=8.5. 8 ul of TMTPro reagent (ACN stock @ 12.5 mg/mL) were added and labeling were allowed to proceed for 30 min @ RT. Excess of label was quenched by adding 40 ul of 500 mM ABC buffer (30 min @ 37oC). Labeled peptides were mixed together to create 2 × 16 plex TMT batches, that were subsequently desalted on tC18 SepPak cartridges, concentrated on speedvac and fractionated offline.

500 ug of peptides were fractionated using a Waters XBridge BEH 130A C18 3.5um 4.63mm ID x 250 mm column on an Agilent 1260 Infinity series HPLC system operating at a flow rate of 1 mL/min with three buffer lines: Buffer A consisting of water, buffer B of ACN and Buffer C of 100 mM ammonium bicarbonate. Peptides were separated by a linear gradient from 5% B to 35% B in 62 min followed by a linear increase to 60% B in 5 min, and ramped to 70% B in 3 min. Buffer C was constantly introduced throughout the gradient at 10%. Fractions were collected every 60 s. Fractions from 30 to 64 were used for LC-MS/MS analysis.

LC separation was performed online on EvosepOne LC1 utilizing Dr Maisch C18 AQ, 1.9μm beads (150μm ID, 15cm long, cat# EV-1106) analytical column. Peptides were gradient eluted from the column directly to Orbitrap HFX mass spectrometer using 44 min evosep method (30SPD) at a flowrate of 220 nl/min. Mass spectrometer was operated in either data-dependent acquisition mode DIA. High resolution full MS spectra were acquired with a resolution of 120,000, an AGC target of 3e6, with a maximum ion injection time of 100 ms, and scan range of 400 to 1600 m/z. Following each full MS scan 20 data-dependent HCD MS/MS scans were acquired at the resolution of 60,000, AGC target of 5e5, maximum ion time of 100 ms, one microscan, 0.4 m/z isolation window, nce of 30, fixed first mass 100 m/z and dynamic exclusion for 45 seconds. Both MS and MS2 spectra were recorded in profile mode.

### Proteome analysis pipeline

MS data were analyzed using MaxQuant software version 1.6.15.02 and searched against the SwissProt subset of the human uniprot database <u>(h</u>t<u>tp://www.uniprot.org/)</u> containing 20,430 entries. Database search was performed in Andromeda3 integrated in MaxQuant environment. A list of 248 common laboratory contaminants included in MaxQuant was also added to the database as well as reversed versions of all sequences. For searching, the enzyme specificity was set to trypsin with the maximum number of missed cleavages set to 2. The precursor mass tolerance was set to 20 ppm for the first search used for non-linear mass re-calibration4 and then to 6 ppm for the main search. Oxidation of methionine was searched as variable modification; carbamidomethylation of cysteines was searched as a fixed modification. TMT labeling was set to lysine residues and N-terminal amino groups, corresponding batch-specific isotopic correction factors were accounted for. The false discovery rate (FDR) for peptide, protein, and site identification was set to 1%, the minimum peptide length was set to 6. To transfer identifications across different runs, the ‘match between runs’ option in MaxQuant was disabled. Only precursors with minimum precursor ion fraction (PIF) of 75% were used for protein quantification. Match between runs option was enabled and RAW TMT reporter ion intensities of peptide features were used for subsequent data analysis in MSstatsTMT5.

Subsequent data analysis were performed in either Perseus6 (http://www.perseus-framework.org/) or using R environment for statistical computing and graphics (http://www.r-project.org/).

### Quantification of aneuploidy degree

The Segment files for different cancer types were from CPTAC. We adjusted the segments to a 100kb window size and the arm-level copy number alterations were calculated based on copy number package (Nilsen et al., 2012). We considered a log2-transformed copy number ratio >0.2 as a gain and <(−0.2) as a loss. The aneuploidy degree corresponds to the total number of chromosome arm gains or losses (any chromosome).

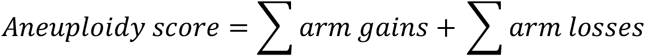

The aneuploidy degree of CCLE was downloaded from geneDep website (Cohen-Sharir et al., 2021)

### Evaluation of the association of gene expression with aneuploidy by linear model

To find the genes whose protein expression changes along with aneuploidy, a linear model was used to fit the protein expression (for cell lines or individual tumor tissues) or the protein or RNA log2FC (for pooled tumor tissues) by aneuploidy score and other covariates including cancer types, copy number variation, purity or cell cycle score. One example is shown as the following formula.

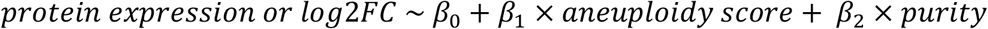

The t-value of aneuploidy coefficient *β*_1_was used to represent the association between protein level and aneuploidy degree with the control of other variables (such as purity). The genes were ranked based on the t-value of aneuploidy coefficient *β*_1_and then the enrichment of gene sets was calculated by Gene Set Enrichment Analysis (GSEA) with preranked module. C5 BP gene sets derived from the GO biological Process ontology were used in those analyses. The gene sets whose size were smaller than 5 or bigger than 500 were removed before analyses.

As gene sets related to transcription and translation include many mitochondrial ribosome, rRNA and tRNA genes, we also removed all mitochondrial genes before GSEA to exclude the possibility that mitochondrial genes overwhelm those gene sets. For purity scores, CPTAC has two sets of purity data from nuclei percentage and the estimated amount of immune infiltrate based on the algorithm Estimate. Data from the algorithm Estimate are missing for ccRCC and OV. For the data from nuclei percentage, COAD, BRCA and OV are missing. Those missing cancers are excluded from the pan-cancer analysis when purity was included in the model. The cell cycle score was calculated based on the average level RNA level of ten genes related to cell cycle entry (Davoli et al., 2017). To compare CPTAC and CCLE, the common genes of those two datasets were used for linear model and GSEA. The genes used to analyze changes at the RNA levels were the same ones used for analysis of protein change.

## Codes Availability

Codes used in this manuscript are available in GitHub, https://github.com/breezyzhao/Proteogenomic-Analysis-of-Aneuploidy. All other study data are included in the article and supplemental information.

## Acknowledgments

We thank all the members of the Davoli and Fenyö labs as well as members of the Kelly Ruggles lab and Christine Vogel (NYU) for helpful comments and insights during the completion of the project.

